# st-DenseViT: A Weakly Supervised Spatiotemporal Vision Transformer for Dense Prediction of Dynamic Brain Networks

**DOI:** 10.1101/2024.11.28.625914

**Authors:** Behnam Kazemivash, Pranav Suresh, Dong Hye Ye, Armin Iraji, Jingyu Liu, Sergey Plis, Peter Kochunov, Vince D. Calhoun

## Abstract

**Objective:** Modeling dynamic neuronal activity within brain networks enables the precise tracking of rapid temporal fluctuations across different brain regions. However, current approaches in computational neuroscience fall short of capturing and representing the spatiotemporal dynamics within each brain network. We developed a novel weakly supervised spatiotemporal dense prediction model capable of generating personalized 4D dynamic brain networks from fMRI data, providing a more granular representation of brain activity over time.

**Methods:** We developed a model that leverages the vision transformer (ViT) as its backbone, jointly encoding spatial and temporal information from fMRI inputs using two different configurations: space-time and sequential encoders. The model generates 4D brain network maps that evolve over time, capturing dynamic changes in both spatial and temporal dimensions. In the absence of ground-truth data, we used spatially constrained windowed independent component analysis (ICA) components derived from fMRI data as weak supervision to guide the training process. The model was evaluated using large-scale resting-state fMRI datasets, and statistical analyses were conducted to assess the effectiveness of the generated dynamic maps using various metrics.

**Results:** Our model effectively produced 4D brain maps that captured both inter-subject and temporal variations, offering a dynamic representation of evolving brain networks. Notably, the model demonstrated the ability to produce smooth maps from noisy priors, effectively denoising the resulting brain dynamics. Additionally, statistically significant differences were observed in the temporally averaged brain maps, as well as in the summation of absolute temporal gradient maps, between patients with schizophrenia and healthy controls. For example, within the Default Mode Network (DMN), significant differences emerged in the temporally averaged space-time configurations, particularly in the thalamus, where healthy controls exhibited higher activity levels compared to subjects with schizophrenia. These findings highlight the model’s potential for differentiating between clinical populations.

**Conclusion:** The proposed spatiotemporal dense prediction model offers an effective approach for generating dynamic brain maps by capturing significant spatiotemporal variations in brain activity. Leveraging weak supervision through ICA components enables the model to learn dynamic patterns without direct ground-truth data, making it a robust and efficient tool for brain mapping.

**Significance:** This work presents an important new approach for dynamic brain mapping, potentially opening up new opportunities for studying brain dynamics within specific networks. By framing the problem as a spatiotemporal dense prediction task in computer vision, we leverage the spatiotemporal ViT architecture combined with weakly supervised learning techniques to efficiently and effectively estimate these maps.

## I. Introduction

The human brain, an incredibly complex system, consists of nearly one hundred billion neurons, each forming tens of thousands of unique connections with other neurons [1]–[3]. This forms complex computational and memory networks that exhibit non-linear dynamics underpinning essential functions such as cognition, memory, emotion, and perception. This dynamic behavior is key to shaping conscious experience and understanding human cognition and consciousness [4]–[6].

Advances in neuroimaging, particularly functional magnetic resonance imaging (fMRI), have significantly transformed computational neuroscience by enabling the simultaneous recording of blood oxygenation level dependence (BOLD) signal activity across the entire brain [7]–[9]. Many analytical and modeling techniques have been developed to investigate brain dynamics using sequential neural data. One widely used method is based on co-activation patterns (CAPs), which identifies transient brain patterns that contribute to the formation of resting-state networks [10], [11]. CAPs provide an effective method for identifying recurring neural configurations but struggle to resolve meaningful patterns when functionally distinct brain states exhibit significant temporal overlap, potentially limiting their sensitivity in capturing complex or concurrent neural processes [12]–[14].

Another prominent approach is sliding window correlation (SWC), commonly used to investigate dynamic changes in regional connectivity. This method involves calculating pairwise correlation or covariance matrices within a specified time window across successive time points [15], [16]. While it is effective for tracking the temporal evolution of functional connectivity, the choice of window length can significantly influence the results. Newer models attempt to relax this constraint [17]. However, overlapping windows may introduce correlated noise and redundancy, complicating the distinction between true dynamic changes and artifacts in the data [18]– [20].

Similarly, phase synchrony (PS) has been used to measure coherence and phase coupling between neural components, providing insights into the synchrony between brain regions [21], [22]. This method is well-suited for capturing the temporal coordination of neural activity, however, it is primarily focused on phase relationships and may not fully capture the amplitude variations [23]–[25]. Switching linear dynamical systems (SLDS) are another modeling approach used to represent nonlinear dynamics by approximating complex systems through transitions between multiple linear regimes [26]. These models are flexible and can represent nonlinear behaviors in brain activity, but the need for multiple discrete states to approximate a single nonlinear vector field can make the models difficult to interpret. The number of latent dimensions required for each regime further complicates the visualization and interpretation of results. [27].

Although many methods focus on temporal variations across fixed spatial nodes, recent work emphasizes the importance of capturing spatial variability within functional domains for a more comprehensive understanding of brain activity [28]. Quasi-periodic patterns (QPPs) capture reliable spatiotemporal patterns in low-frequency neural activity [29], [30]. This approach is valuable for uncovering recurring dynamics over time, offering insights into both temporal and spatial interactions in brain function. However, QPPs are primarily focused on low-frequency activity, potentially limiting their ability to detect higher-frequency events critical for understanding brain dynamics [31]–[33]. Hierarchical models that integrate high-order spatial independent component analysis (ICA) reveal the fluidity of spatial associations, distinguishing functional homogeneity and stability at lower levels from greater spatial variability at higher levels [34], [35]. Windowed ICA approaches provide even more flexibility, allowing spatially varying maps for each brain network [34].

Although each method offers distinct advantages, a comprehensive understanding of brain dynamics requires analyzing both the temporal evolution of individual networks and the interactions between different brain regions over short timescales. The generation of dynamic brain networks can be reframed as a computer vision problem, using UNet-style models to capture spatially evolving patterns in brain data [36], [37]. These generated maps have been effective in distinguishing between schizophrenia and control groups, demonstrating their potential for identifying subtle alterations in brain activity [38], [39]. While these innovative methods advance the creation of dynamic brain maps, the development of robust parcellation techniques that yield detailed, temporally resolved maps remains a critical challenge in the field.

In this study, we present a novel model that addresses three challenges in analysis of dynamic brain networks. First, the model introduces a soft brain parcellation technique that produces high-resolution 4D brain maps. Second, it functions as a dynamic estimation tool, capturing and representing distinct spatiotemporal activity patterns across multiple brain networks over time. Third, it overcomes the absence of ground-truth data by incorporating spatially constrained windowed ICA components [40] as weak supervision to initialize the training. Finally, we assess the medical relevance and performance of the generated 4D maps to highlight the practical impact of our approach.

## II. Related Concepts

In this section, we explore key concepts central to our approach, including weakly supervised learning, spatiotemporal dense prediction, and brain parcellation. Understanding these foundational ideas is essential for grasping the methods and objectives of our research on dynamic brain mapping.

### A. Weakly Supervised Learning

Weakly supervised learning approaches allow predictive models to be trained using datasets that lack precise, fully labeled examples. Instead of relying on exact labels for every data point, these methods can handle noisy, ambiguous, or incomplete data [41], [42]. One common scenario in weakly supervised learning is label uncertainty, where each training instance is associated with a distribution of possible labels rather than a single definitive label. In such cases, training is performed by minimizing the cross-entropy between the true label distribution and the model’s predicted distribution, a technique known as label smoothing. This method can regularize the model by replacing hard labels with softer, probabilistic versions [43]–[45].

Another method in weakly supervised learning is multiple instance learning (MIL). In MIL, training data is grouped into sets or “bags” where only a label for the entire bag is available, but not for individual instances within the bag [46]–[48]. Distant supervision is yet another strategy in this framework, where labels are inferred from external knowledge sources, such as databases [49]–[51]. Label-noise learning (LNL), which addresses issues caused by incorrect or noisy labels from sources like human error, data uncertainty, or subjective labeling criteria, along with other methods, enables machine learning models to be trained effectively even when ground-truth data is sparse, incomplete, or noisy [52]–[55]. In the era of big data, where obtaining fully labeled datasets is often impractical, weakly supervised learning has become an essential approach.

### B. Spatiotemporal Dense Prediction

In computer vision, dense prediction is a task of pixel-wise or voxel-wise predictions, where the model generates outputs at the level of individual picture or volume elements, rather than at higher levels of abstraction [56]–[58]. Examples of dense prediction tasks include semantic [59], [60], panoptic [61], [62], and instance segmentation [63], [64], optical flow estimation [65], [66], depth estimation [67], [68], and even image reconstruction [69], [70].

The dense prediction models need to accurately capture both spatial and temporal information to produce coherent spatiotemporal representations [71], [72]. To address these challenges, recent techniques have been developed that aggregate flow-guided features, leverage sequence modeling, and apply heuristic approaches to model temporal variations. These methods aim to exploit redundancies and correlations across different time points, improving both predictive accuracy and efficiency [73].

### C. Brain Parcellation

Brain parcellation refers to the segmentation of the brain into distinct regions that are assumed to be functionally or structurally distinct [74]. The simplest approaches are atlas-based methods that rely on predefined anatomical templates with rigid boundaries (i.e., hard parcellation) and therefore face limitations due to individual variability in brain size, shape, and folding, as well as the computational demands of spatial registration [75], [76].

Alternatively, segmentation approaches use functional connectivity [77] parcellation that clusters voxels based on their connectivity profiles, providing an individual-level data-driven representation of brain organization [78]. Among connectivity-based approaches, independent component analysis (ICA), a soft parcellation technique (i.e., allowing voxels to contribute to multiple networks with varying weights), is widely employed [79]. ICA extracts independent spatial maps and captures the temporal contribution of these maps through corresponding time courses. However, reducing these temporal contributions to scalar values per volume may oversimplify the underlying complexity of brain activity [28], [80], [81]. Consequently, there remains a critical gap in the development of advanced dynamic brain parcellation methods capable of fully capturing the intricate temporal dynamics present in neural data and generating spatiotemporally evolving brain activity patterns that accurately reflect the continuous nature of functional brain processes.

### III. Methods

Our model generates dynamic patterns of brain networks by employing two distinct configurations independently: a space-time encoder and a sequential encoder, as shown in Figure 1. Additionally, we propose a method to tackle the challenge of missing ground-truth data when generating dynamic brain maps.

**Fig. 1.**
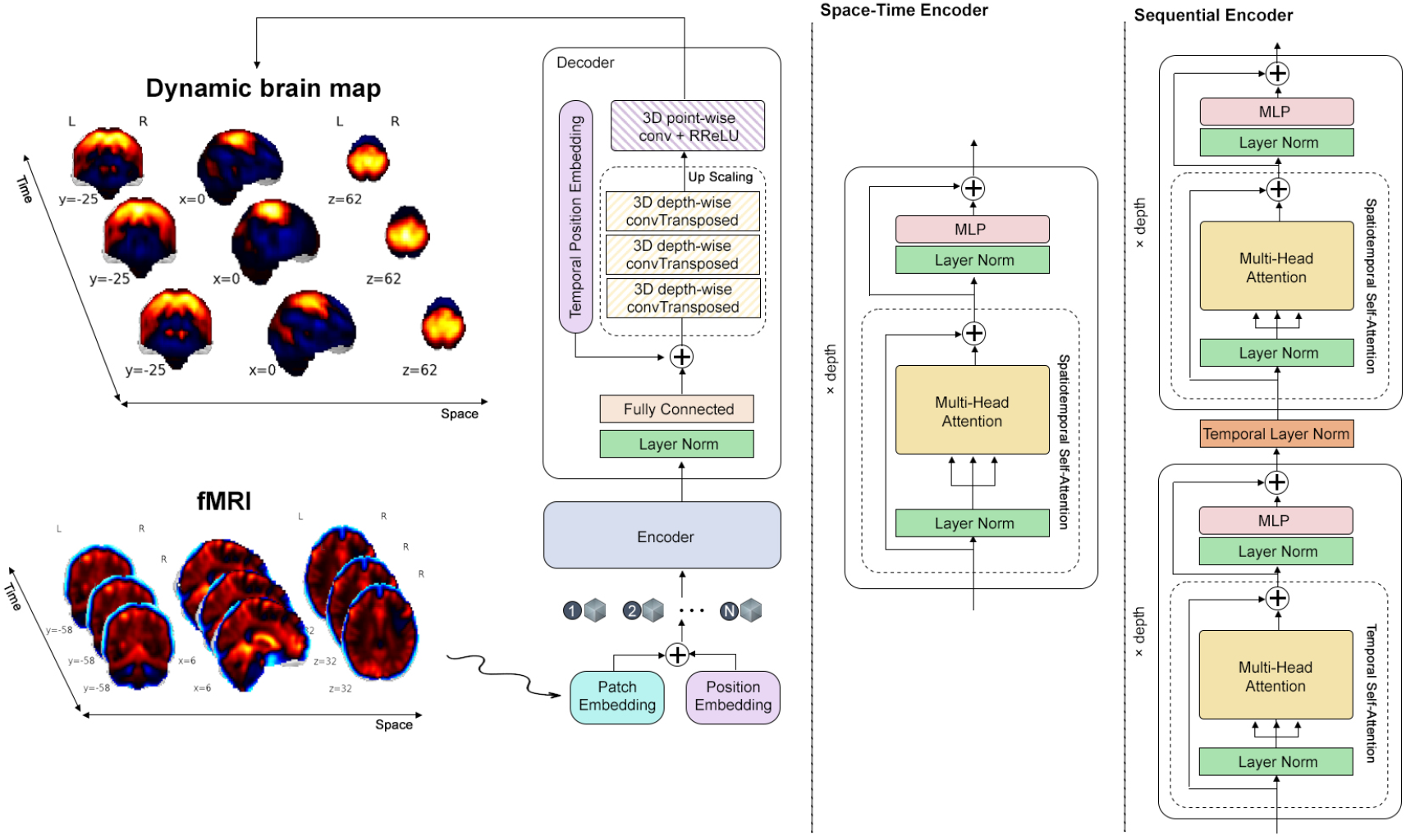
We present a schematic representation of our proposed spatiotemporal dense prediction model, aimed at generating dynamic brain maps. To effectively capture dynamic patterns, we employ a vanilla Vision Transformer (ViT) encoder that processes a substantial number of spatiotemporal tokens. Notably, we have developed two variants of the encoder: a space-time encoder and a sequential encoder, both designed to facilitate the learning of spatiotemporal information. Additionally, as illustrated in the figure, we utilize a CNN decoder head to reconstruct the extracted features into the original spatial dimensions.

We utilize a vision transformer as the backbone of our model. First, we patchify the input fMRI data, *x* ∈ ℝ^*h×w×d×t*^, into a sequence of tokens 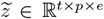. More precisely, we extract *p* non-overlapping 3D patches *s*_*i*_ from the spatial dimensions, apply a linear projection *π*, and then flatten them into 1D tokens *z*_*i*_ ∈ ℝ^*e*^. Additionally, a learnable positional embedding, *pe* ∈ ℝ^*t×p×e*^, is added to the tokens to retain positional information, addressing the permutation invariance of self-attention module within the transformer. So we have 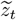 representing a token for a timepoint *t* as follows:

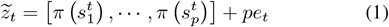

### A. Space-Time Configuration

In our first scenario, the encoder architecture is designed to process spatial tokens across all time points, effectively capturing both spatial and temporal information. The encoder consists of a stack of *L* transformers, each comprising modules such as multi-headed self-attention, layer normalization, and a multi-layer perceptron (MLP) with two fully connected layers, dropout, and randomized leaky rectified linear unit function (RReLU) activation, as outlined below:

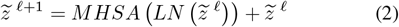

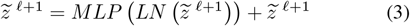

Furthermore, the attention mechanism is defined as follows, where the queries *Q*, keys *K*, and values *V* are linear projections of the input, with *Q, K, V* ∈ ℝ^*p×e*^, and *e* representing the embedding dimension. All spatiotemporal tokens extracted from input data are then passed through the transformer encoder to capture dynamic patterns:

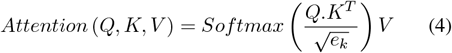

### B. Sequential Encoders Configuration

In our second scenario, we employ a ViT with two distinct modules for temporal and spatial encoding to process temporal and spatial tokens separately. First, we permute the tensor of token sequences, 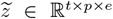, to obtain a new shape 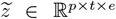. This permuted tensor is then fed into the temporal encoder. Afterward, we reshape the output back to the original shape 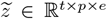 and feed it into the spatial encoder. The embedding dimensions of both the temporal and spatial encoders are identical.

### C. Decoder Head

Unlike the encoder, which is built on the ViT architecture, the decoder head is designed with a series of components. It consists of a layer normalization, followed by a fully connected layer, a fixed sine-cosine positional encoding to represent the time-point indices, and a sequence of 3D depthwise transposed convolution operator (ConvTranspose) layers with various kernel sizes of 7, 5, and 9, respectively. The final ConvTranspose layer has a dilation rate of 2. The decoder also includes a point-wise Conv3D layer, with RReLU serving as the activation function.

### D. Customized Loss function

Selecting an appropriate loss function that simultaneously preserves the global structure of the data while capturing dynamic patterns is a challenging task. To address this, we utilize a combination of photometric and perceptual losses to guide the network’s training, as formulated below:

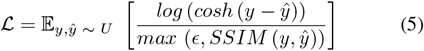

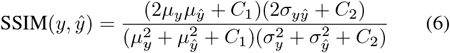

Here, *y* represents the prior (weak supervision), and *ŷ* corresponds to the model’s prediction, while SSIM denotes the structural similarity index. The parameter *ϵ* = 0.001 is used to prevent division by zero or negative values in the denominator and *C*1, *C*2 are small constants to stabilize the division. The *log*(*cosh*(.)) function serves as a robust regression loss, similar to mean squared error (MSE), but less prone to being influenced by outliers. It offers the benefits of the Huber loss while remaining continuously differentiable, unlike the Huber loss. Additionally, we observe empirically that incorporating SSIM loss creates more opportunities for improving prediction quality.

### E. Weak Supervision

One of the main challenges in our research is the lack of ground-truth labels, which we addressed by utilizing a windowed version of spatially constrained ICA [40], a powerful semi-blind source separation technique [82]. This method enables us to extract independent components (ICs), each representing a distinct brain network, which we used as weak supervision to guide the training of our model.

Our method leverages constrained ICA by using the NeuroMark template [83], which consists of reproducible independent components derived from the Genomic Superstructure Project (GSP) and Human Connectome Project (HCP) datasets. These group-level constrained windowed ICs were computed by spatially aligning the correlated components from more than 800 healthy control fMRI datasets. The NeuroMark template provides a reference for calculating subject-specific ICs using a multi-objective optimization strategy as follows:

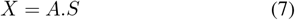

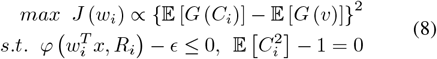

In this context, *X* denotes the input fMRI data, *A* is the unknown mixing matrix, and *S* represents the independent sources. The term *J* (*w*_*i*_) estimates the negentropy of the *i*th source *C*_*i*_, with *w*_*i*_ as its corresponding unmixing matrix. 𝔼 [.] stands for the expectation, while *G* (.) is a non-quadratic function. The variable *v* is a Gaussian random variable with a mean of zero and unit variance. The function *φ* measures the distance between the estimated source *C*_*i*_ and its normalized reference *R*_*i*_, with *ϵ* acting as a threshold to control the separation accuracy between the independent component and the reference signal. We utilize spatially constrained ICA on fMRI data by implementing overlapping windows, with a window size of 30 and a stride of 1, to extract a sequence of independent components. This approach offers advantages in extracting brain networks compared to other methods, though it remains a linear technique. Additionally, this method is sensitive to noise, and in cases of low signal-to-noise ratio, the accuracy of component separation may be affected. In the windowed ICA configuration, it is also assumed that the statistical properties of the sources remain constant, an assumption that may not always hold. Despite these limitations, spatially constrained windowed ICA can serve as a suitable weak prior for a deep learning model, as we propose in this work. A key strength of our approach is its ability to capture nonlinearities in the spatial domain of brain networks.

## IV. Experiment

We implemented a straightforward experimental setup for our research. Our study utilized a subset of 508 fMRI datasets, which included data from the MPRC [84], FBIRN [85], and COBRE [86] datasets. The demographic information for this subset is outlined in Table I. To reduce computational demands, we uniformly sampled time points from the fMRI data at 10-timepoint intervals, resulting in a total of 10 time points instead of using the full dataset. We also applied a Gaussian filter for image smoothing, with a standard deviation of *σ* = 6 set for Gaussian kernel, followed by z-scoring to ensure proper normalization of the data. The model configuration included an embedding dimension of 96, 6 attention heads, a depth of 1, an attention dropout rate of 0.4, and an encoder dropout rate of 0.3, with a patch size of 5. Training was conducted on two A40 GPUs, each equipped with 45 GB of memory, using a batch size of 2. We utilized the Adam optimizer with a learning rate of 0.01 and a weight decay of 0.1, running for a total of 150 epochs and implementing an early stopping strategy to prevent overfitting, with a threshold of *δ* = 10^−5^.

**TABLE 1.** Demographic information.

| Diagnostic | Sex |  | Age<br>mean+/-sd | Race |  |  |
| --- | --- | --- | --- | --- | --- | --- |
|  | Male | Female |  | American | European | Other |
| Control | 185 | 130 | 38.40 +/- 12.73 | 192 | 65 | 58 |
| Schizophrenia | 154 | 39 | 38.61 +/- 13.29 | 112 | 44 | 37 |

### A. Qualitative Metrics

We present qualitative comparisons between the dense visual representations learned by our model and the extracted sequence of ICA components (used for weak supervision) across different networks, employing both space-time and sequential encoder configurations, as shown in Figure 2. The dynamic patterns of brain activities observed in the generated maps exhibit smooth transitions in both space and time, with noticeable fluctuations in activation weights (scores) across different time points. These transitions highlight the model’s capability to capture temporal continuity, resulting in more plausible representations of brain activity. The generated maps show a strong alignment with known brain atlases, particularly in the spatial localization of functional regions, reinforcing the model’s accuracy in learning meaningful spatial features. Moreover, we provide additional evidence of the model’s ability to capture dynamic patterns over time by calculating the summation of absolute temporal gradients, approximated using the forward difference approach. This metric quantifies the temporal changes in brain activity, offering a complementary view of how the model captures both gradual and abrupt transitions across time points. The temporal gradients underscore the model’s sensitivity to changes in activation dynamics, further validating its capacity to generate temporally consistent and biologically meaningful representations.

**Fig. 2.**
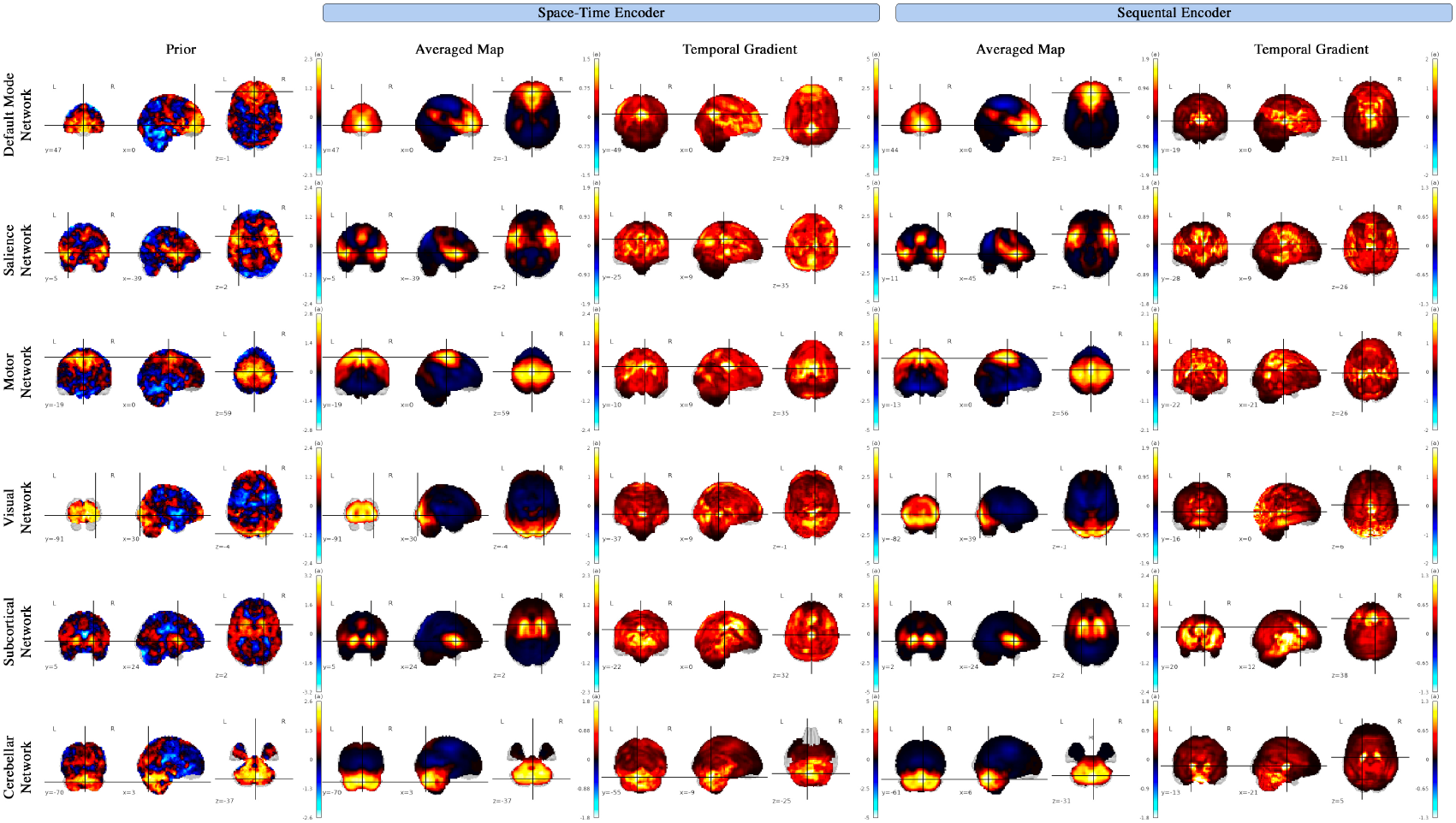
Model-generated dynamic brain maps within the Default Mode, Salience, Motor, Visual, Subcortical, and Cerebellar networks, averaged over time for a randomly selected test subject. The figure includes the summation of absolute temporal gradient maps and prior maps, comparing outputs from both the Space-Time encoder and Sequential encoder configurations.

Additionally, the subject-specific variability in a sample cerebellar network for a sample timepoint 5, as illustrated in Figure 3, emphasizes the model’s strength in adapting to individual differences in brain dynamics, revealing subtle but significant deviations from population-level patterns. This variability is crucial for understanding the heterogeneity in brain function across different individuals. Furthermore, the model’s robustness is evidenced by its ability to effectively denoise highly noisy prior (weak supervision), refining them into clearer, more interpretable dynamic brain maps. This denoising capability not only improves the signal-to-noise ratio but also enhances the interpretability of the spatiotemporal patterns, facilitating a deeper understanding of the underlying neural processes. Overall, the model excels in translating raw, noisy data into coherent, high-resolution representations that capture both the global and individualized aspects of brain activity over time within each brain network.

**Fig. 3.**
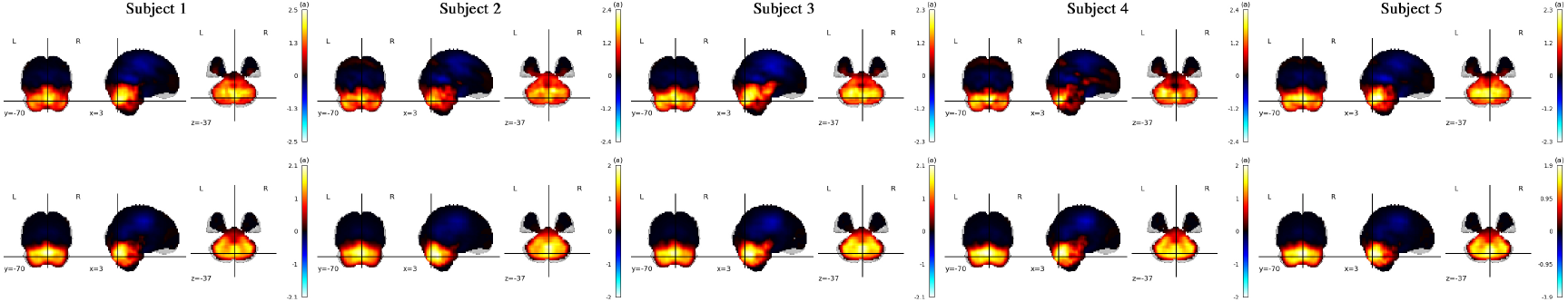
Generated maps for five distinct subjects at the fifth time point reveal substantial inter-subject variability, highlighted by differences in morphology, scale, spatial displacement, and activation intensity of active regions. The top row displays the space-time configuration, while the bottom row illustrates the sequential encoder configuration.

To assess the clinical relevance of the generated dynamic brain maps, we conducted a series of experiments to explore their potential for differentiating between healthy controls (HC) and individuals with schizophrenia (SCZ). The dynamic brain maps, generated by our model for all subjects, included both HC and SCZ groups. In the first experiment, we averaged the dynamic maps across time and applied a mask to extract brain voxels, *HC* ∈ ℝ^315*×*68235^ and *SCZ* ∈ ℝ^193*×*68235^. A voxel-wise t-test was then performed to identify regions with significant differences (*p* ≤ 0.05) between the two groups. This analysis enabled the detection of specific brain regions that exhibit altered dynamics in SCZ, potentially revealing biomarkers for the disorder. In the second experiment, we applied the same t-test to the TG maps, which capture voxelwise differences between consecutive time-points, to assess group differences in dynamic brain activity. This approach allows us to study variations in dynamic brain patterns, as shown in Figure 4, and their potential association with the pathophysiology of schizophrenia. These findings lay a foundation for further exploration of their relevance in clinical observations and future research on diagnostic and therapeutic strategies.

**Fig. 4.**
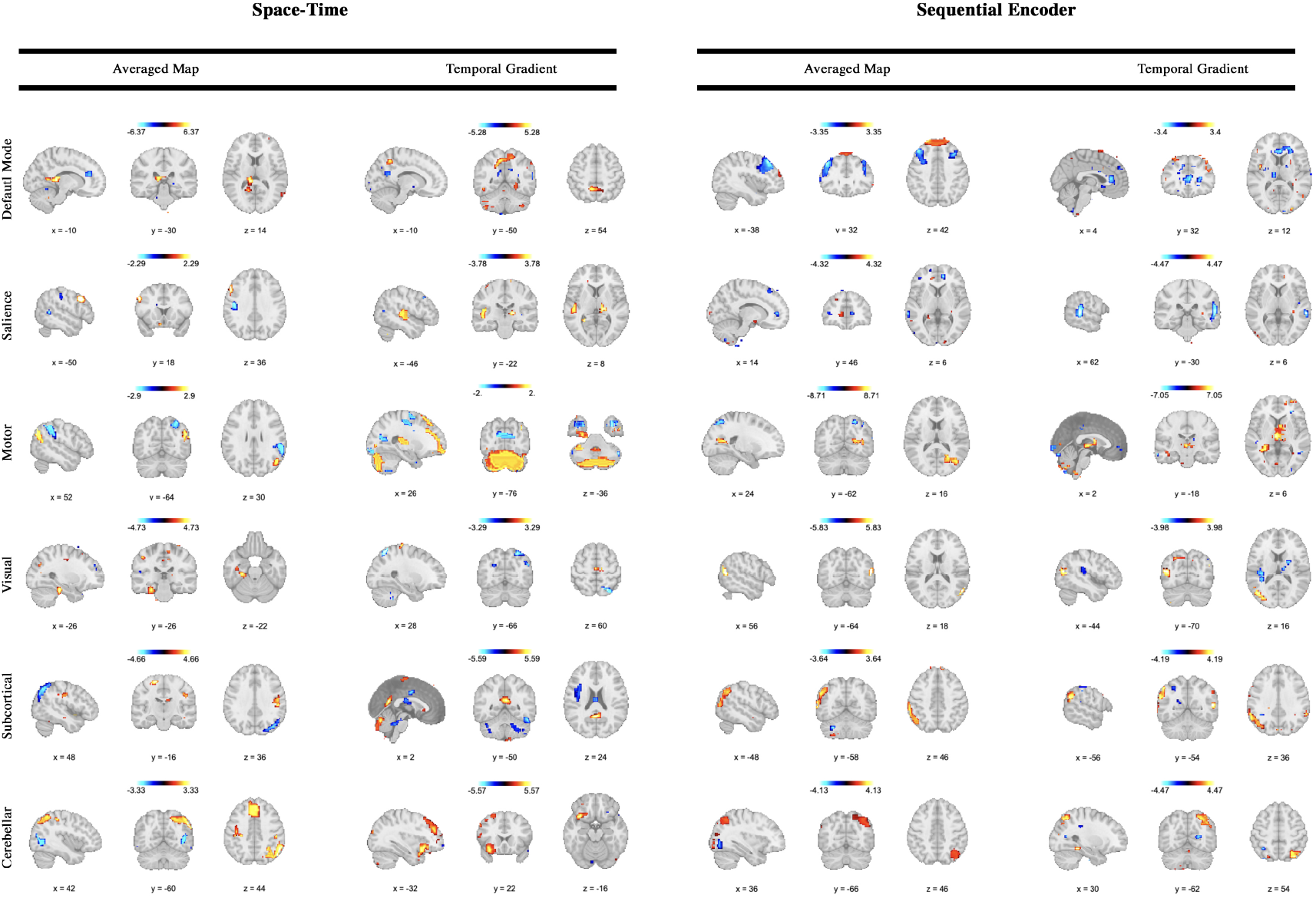
Group-level differences across multiple regions are highlighted in both the averaged maps and the sum of temporal gradients, assessed using a two-sample t-test with FDR correction and masked at ***p* ≤ 0.05** across the default mode, salience, motor, visual, subcortical, and cerebellar networks. The maps display **− log(*p _value*) *×*** sign**(*t_value*)**, emphasizing the relevance of the generated dynamic maps to our understanding of schizophrenia. Widespread group differences are observed across several brain regions, with schizophrenia associated with both spatial and temporal changes. While healthy controls (HCs) generally exhibit greater hyperactivity and variations, schizophrenia demonstrates increased activity and variation in specific regions. Notably, these changes often exhibit structured patterns that do not strictly align with regions strongly contributing to the networks, suggesting the presence of transient states or functional connectivity across networks. The left two columns illustrate the space-time configuration, whereas the right two columns present the sequential encoder configuration, highlighting differences between the two scenarios.

### B. Quantitative Metrics

We quantitatively assess the model’s capacity to generate plausible dense representations using an indirect standard approach. This evaluation focuses on several key metrics, including active region localization accuracy, visual fidelity, consistency with expected patterns, and regional homogeneity, which is quantified by the correlation of voxel time-series within each region of interest (ROI). We perform these assessments on both the generated maps and the priors, averaging results over time, while applying a threshold of ≥ 80% specifically to the mean intersection over union (mIOU) and homogeneity (Hgt) metrics, as detailed in Table II. Additionally, we clarify that mARE denotes the mean absolute relative error, mIOU refers to the mean intersection over union, and SSIM signifies the structural similarity index.

**TABLE II.** Quantitative metrics on quality of generated dynamic maps.

|  | *Networks* | *mARE* ↓ | *mIOU* ↑ | *SSIM* ↑ | *Hgt* ↑ |
| --- | --- | --- | --- | --- | --- |
| Space-Time | Default Mode | 0.12 | **0.77** | **0.87** | 0.77 |
|  | Salience | 0.17 | 0.71 | 0.79 | 0.79 |
|  | Motor | 0.14 | 0.73 | 0.86 | 0.77 |
|  | Visual | 0.15 | **0.77** | 0.84 | 0.81 |
|  | Subcortical | 0.15 | **0.77** | 0.81 | 0.72 |
|  | Cerebellar | **0.1** | 0.81 | 0.86 | **0.84** |
| Sequential Enc | Default Mode | 0.15 | 0.73 | **0.85** | 0.7 |
|  | Salience | 0.21 | 0.69 | 0.73 | 0.63 |
|  | Motor | 0.17 | 0.72 | 0.8 | 0.74 |
|  | Visual | 0.16 | 0.77 | 0.8 | 0.72 |
|  | Subcortical | 0.17 | 0.74 | 0.75 | 0.66 |
|  | Cerebellar | **0.13** | **0.79** | 0.82 | **0.79** |

The quantitative results reveal several interesting patterns across networks and encoding schemes. The Cerebellar network stands out with the best overall performance, achieving the lowest mARE and highest Hgt in both Space-Time (mARE: 0.10, Hgt: 0.84) and Sequential Enc (mARE: 0.13, Hgt: 0.79), along with competitive mIOU and SSIM scores. In contrast, the Salience network consistently underperforms, with the highest mARE and lowest mIOU, SSIM, and Hgt in both encoding schemes, suggesting challenges in capturing its diffuse and variable activity patterns. The Default Mode network excels in structural similarity, achieving the highest SSIM (0.87 in Space-Time and 0.85 in Sequential Enc), indicating robust reconstruction of its coherent activity patterns. The Visual network demonstrates stable performance, with relatively high mIOU and Hgt scores across both schemes, reflecting the predictable nature of visual dynamics. Notably, Space-Time encoding generally results in lower mARE values, highlighting better reconstruction accuracy, while Sequential Enc slightly outperforms in mIOU for the Cerebellar and Subcortical networks, suggesting improved spatial overlap capture in specific cases. These findings are consistent with the expected dynamic characteristics of these brain networks.

Furthermore, we evaluate the model’s capacity to generate dynamic patterns over time by calculating the Shannon entropy of the temporal gradient maps, capturing changes in information content as an additional indication of the model’s dynamic behavior. This metric enables quantification of evolving patterns across temporal dimensions, enhancing our understanding of the underlying brain dynamics, as illustrated in upper row of Figure 5.

**Fig. 5.**
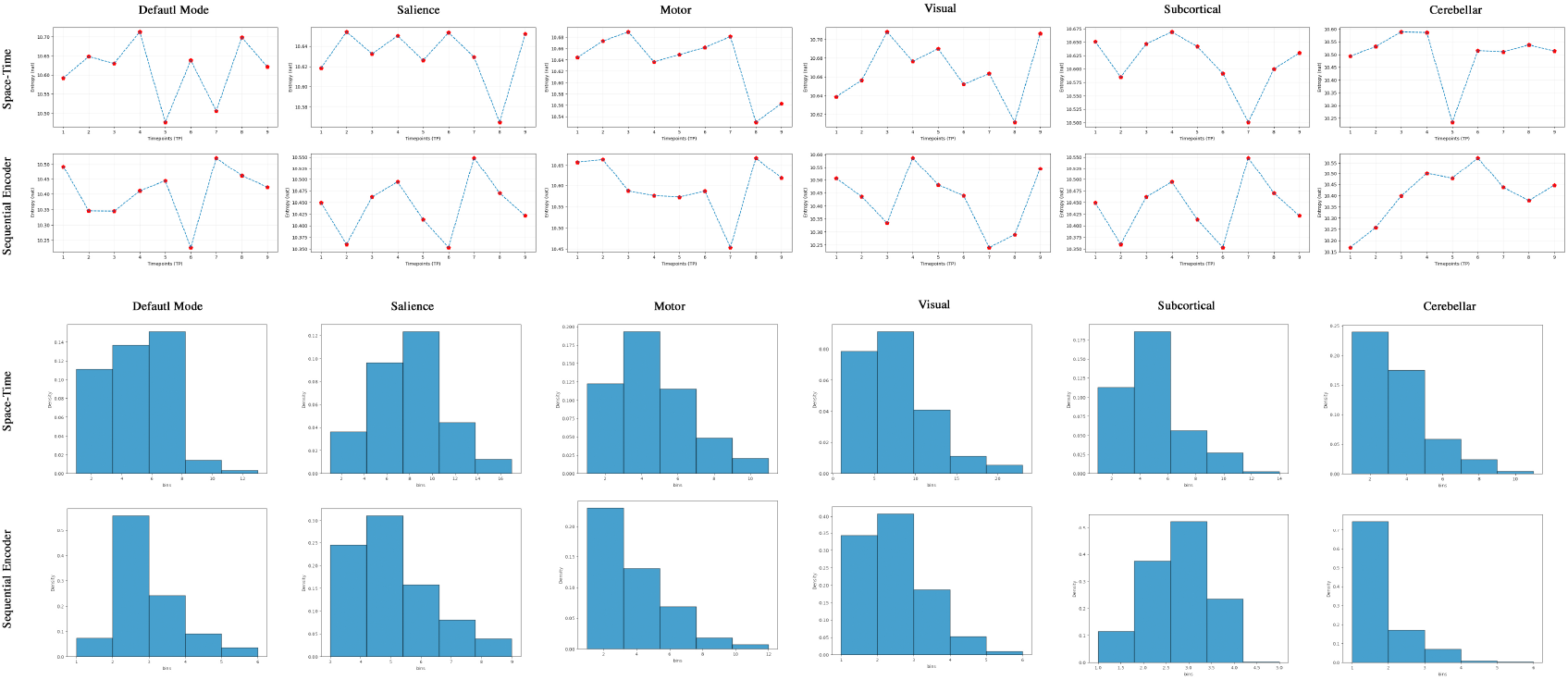
Shannon entropy trends (computed for a single random test subject) and blob density histograms (calculated across the entire test set) for six brain networks: default mode, salience, motor, visual, subcortical, and cerebellar. These are presented for two encoding schemes: a space-time configuration (rows 1 and 3) and a sequential encoder configuration (rows 2 and 4). The space-time configuration shows a wider range of entropy fluctuations for all networks, indicating greater sensitivity to temporal changes, while the sequential encoder configuration exhibits more stable temporal gradients in Shannon entropy, reflecting consistent representations of spatiotemporal dynamics (e.g., DMN entropy ranges from [10.5, 10.7] for space-time and [10.25, 10.50] for sequential encoder). The blob density histograms reveal distinct behaviors across networks in both configurations. For instance, in the sequential encoder scenario, the cerebellar network shows a tighter and more compact distribution, which is also shifted to lower values compared to other networks, suggesting higher spatial consistency across the test set—most subjects exhibit a similar, smaller number of blobs throughout the entire time interval. In contrast, the space-time configuration displays a wider distribution for the cerebellar network, indicating higher variability while also being shifted toward lower values. Additionally, the motor network in the sequential encoder scenario demonstrates a noticeable shift toward lower values, highlighting reduced activity or variation in blob density for this configuration.

Additionally, we utilize the connected components algorithm as an effective method to measure the dynamicity of the generated 4D maps. By applying this algorithm to the thresholded maps at *δ* = 0.65, we identify blobs at each time point to assess the dynamics. We then track the number of blobs over different networks for the whole test set. This approach is illustrated in lower part of Figure 5. The findings clearly indicate that the active region (ROI) gradually shrinks and merges in the time interval, as evidenced by the changes in the number of blobs. This observation underscores the proposed model’s effectiveness in capturing underlying dynamics and generate dynamic patterns.

## V. Discussion

Our study presents a new approach to capturing brain dynamics that reveal spatiotemporal variations in shape, size, and regional location of active regions. The averaged maps align with the results from established ICA-based models and existing brain atlases [79], [87]. Furthermore our model helps to interpret the outcomes by generating denoised maps. Our quantitative metrics, including a low mean mARE and high values for mIOU, SSIM, and homogeneity (shown in Table II), provide strong evidence of the model’s ability to produce plausible maps that preserve essential spatial patterns. In the absence of ground-truth data, we evaluated the method from multiple perspectives, recognizing that different configurations, such as the space-time and sequential encoders, may capture distinct aspects of the underlying dynamics in the fMRI data. The sequential encoder emphasizes temporal patterns, revealing intriguing new properties of the temporal dynamics, as explored in the clinical application section. On the other hand, the space-time encoder demonstrates stronger quantitative performance, effectively capturing both spatial and temporal dynamics. Together, these configurations provide complementary insights, highlighting the multifaceted nature of brain activity and the value of leveraging multiple perspectives to study its complexity.

Furthermore, the summation of absolute temporal gradient maps (TGs) showcases the model’s ability to capture dynamic brain activity, providing insights that extend beyond those available from averaged maps and aligning well with recent findings in computational neuroscience. For instance, elevated variation is detected in the posterior cingulate cortex (PCC) of the default mode network (DMN), a region invisible in averaged map but known for its role in default mode processing [88]. Similarly, the temporal gradient map for the sequential configuration highlights prominent activity in the thalamus, —a region intricately connected with the medial prefrontal cortex (mPFC) and functionally conserved across mammalian species, including humans [89], [90].

Within the salience network, the temporal gradient map for the space-time configuration highlights significant activity in the PCC, a region implicated in the processing of salient events and facial recognition [91]. In the motor network, the model captures the expected elevated activity in the midbrain [92]. Additionally, the sequential encoder configuration reveals prominent activity in the prefrontal cortex, a central area for motor planning, decision-making, and cognitive control [93]. These observations reflect the interconnected nature of these networks as documented in existing research.

In examining the visual network, the model detects activity in the hippocampus, which aligns with literature suggesting the hippocampus’s role in spatial processing and its functional coupling with visual cortical neurons during natural behavior [94], [95]. In contrast, the sequential encoder configuration highlights dynamic patterns in the thalamus, particularly within the lateral geniculate nucleus (LGN), —a crucial relay center for transmitting visual information from the retina to the primary visual cortex [96], [97].

In the subcortical network, the sequential encoder configuration’s temporal gradient reveals notable variability in the anterior cingulate cortex, a region that serves as the origin of the anterior cingulate-subcortical circuit, providing input to subcortical structures such as the ventral striatum and other related areas [98], [99]. Additionally, we observe pronounced variability in the activity scores of the cerebellum in the cerebellar network under the space-time configuration, whereas the sequential encoder configuration shows greater variability in the basal ganglia. The interconnection between these regions is well-documented, linking the motor and non-motor domains of one subcortical system to the corresponding domain in the other. This anatomical connection supports the idea that cerebellar output can influence the input stage of the basal ganglia and vice versa [100], [101].

Figures 5 provides further quantitative evidence of the model’s ability to capture dynamic brain patterns. In Figure 5, Shannon entropy values are computed over temporal gradient maps for a random test subject across all networks, including the default mode, salience, motor, visual, subcortical, and cerebellar networks. This entropy analysis highlights the model’s sensitivity to variations in temporal dynamics, with the top row representing the space-time configuration and the bottom row the sequential encoder configuration. Higher entropy values across networks indicate the model’s capacity to capture complex, fluctuating activity patterns across these functional regions. Figure 5 further explores dynamic brain patterns through changes in blob count in the time interval. These metrics emphasize fluctuations in activity within the 4D maps, providing insights into the spatiotemporal variation captured by each configuration. Notably, the space-time configuration exhibits more pronounced changes in blob characteristics than the sequential encoder configuration, underscoring its enhanced ability to capture and represent dynamic spatial features in brain activity.

Overall, these findings demonstrate the model’s robustness in capturing complex, dynamic activity patterns across multiple brain networks, highlighting its potential as a tool for examining the temporal and spatial organization of brain function.

### A. Application to Clinical Dataset

In the DMN, there are indications of potential differences in the temporally averaged space-time configurations, particularly within the thalamus, where healthy controls may exhibit higher activity compared to individuals with schizophrenia. This observation aligns, in part, with studies suggesting that disruptions in information flow to and from the thalamus could contribute to symptoms of schizophrenia [34], [102], [103]. Similarly, the TG map suggests possible differences in the parietal lobules, which could indicate dysconnectivity between the parietal lobe and other brain regions in the disorganization symptoms of schizophrenia [104]. Increased activity in the middle frontal gyrus is also tentatively observed in schizophrenia subjects, potentially supporting hypotheses regarding alterations in this region in the condition [105], [106]. Finally, variability in the anterior cingulate cortex (ACC) appears elevated in schizophrenia, resonating with findings that report both hypoactivation and hyperactivation of the ACC. This variability might reflect a hyperactive ACC at rest, which could struggle to further activate under increased task demands, resulting in relative hypoactivity [107].

In the salience network, temporally averaged space-time maps show a tendency for higher prefrontal cortex activation in healthy controls compared to schizophrenia subjects. This observation is partially consistent with research suggesting that schizophrenia patients often face difficulties with tasks dependent on prefrontal cortical function, such as Continuous Performance (attention), Wisconsin Card Sort (cognitive flexibility), Delayed Response (working memory), and N-Back (working memory) tasks [108]. Additionally, there are hints of greater variability in the left primary auditory cortex (Heschl’s gyrus) in healthy controls, which could align with findings linking reduced volume and thickness in this region to auditory hallucinations in schizophrenia [109], [110]. Schizophrenia subjects also seem to show localized activation in a small region of the anterior cingulate and increased variability in the right superior temporal sulcus (STS), a region previously associated with hyperactivation in schizophrenia [111].

In the motor network, temporally averaged maps suggest hyperactivation in the right lateral parietal region of the DMN in healthy controls, while schizophrenia subjects show hyperactivation in the right inferior parietal cortex. These observations may correspond with existing evidence of altered connectivity patterns in schizophrenia, where the lateral DMN exhibits decreased connectivity with sensorimotor regions but increased connectivity with heteromodal association areas [112]. Furthermore, higher variability in cerebellar interactions with motor networks is observed in healthy controls, which could align with studies suggesting disrupted functional associations between cerebellar lobules and motor regions in schizophrenia [113]. Schizophrenia also appears to be associated with hypoactivation in the precuneus [114] and reduced variability in the thalamus compared to healthy controls.

In the visual network, temporally averaged maps suggest hyperactivation in the posterior hippocampus in healthy controls, aligning with previous findings [115]. However, schizophrenia subjects show increased variability in the superior parietal cortex, a region involved in spatial perception, attention, and self-awareness [116]. Similar to the motor network, hypoactivation of the precuneus is observed in schizophrenia. Additionally, reduced variability is seen in the extrastriate cortex for schizophrenia subjects, a region associated with visual processing abnormalities in the disorder [117].

In the subcortical network, individuals with schizophrenia show hypoactivation in the primary motor cortex, which may be linked to motor symptoms arising from disrupted neural circuitry and dysregulated dopamine signaling [118]. However, they also exhibit hyperactivation in a region of the parietal lobe. Furthermore, increased variability is seen in the default mode network in healthy controls, while these controls show hyperactivation in the left parietal lobe and greater variability in the angular gyrus under sequential encoder configurations. Finally, the temporally averaged maps of space-time configuration suggest hyperactivation in the parietal lobes and medial frontal gyrus, regions that are known to be functionally connected [119], [120]. Healthy controls also show greater variability in the orbitofrontal cortex [121]. Sequential encoder configurations indicate hyperactivation in the parietal cortex and heightened variability in the intraparietal cortex in healthy controls compared to schizophrenia subjects. While these trends are intriguing, further investigation is required to better understand the complexities of these network alterations in schizophrenia.

### B. Ablation Study

To evaluate the influence of key design decisions in our model in both space-time and sequential encoder configurations, we performed comprehensive ablation studies focused on two critical aspects: the number of tokens and the temporal resolution. These were systematically varied by modifying patch sizes and the number of time points included in the input data. By experimenting with different patch configurations and temporal granularity, we sought to understand how these factors impact the quality of the results. Specifically, we trained separate model instances for each configuration and assessed their performance on the test set using structural similarity index (SSIM) and mean relative absolute error (mRAE) as evaluation metrics. The findings from these experiments, detailed in Table III and IV, provide valuable insights into the trade-offs associated with tokenization strategies and temporal encoding choices, guiding the optimization of our model’s architecture for enhanced accuracy and robustness.

**TABLE III.**
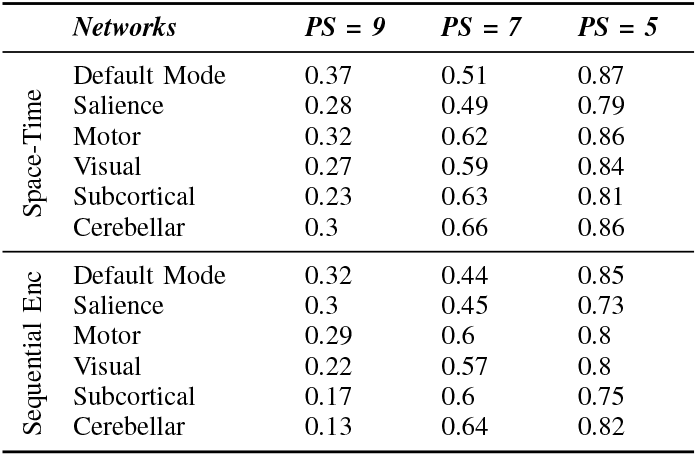
Ablation study on visual quality of generated maps using SSIM.

**TABLE IV.** Ablation study on dynamicity of generated maps using mRAE.

|  | *Networks* | *TP = 10* | *TP = 7* | *TP = 3* |
| --- | --- | --- | --- | --- |
| Space-Time | Default Mode | 0.12 | 0.1 | 0.08 |
|  | Saliency | 0.17 | 0.14 | 0.1 |
|  | Motor | 0.14 | 0.11 | 0.09 |
|  | Visual | 0.15 | 0.13 | 0.1 |
|  | Subcortical | 0.15 | 0.12 | 0.1 |
|  | Cerebellar | 0.1 | 0.08 | 0.05 |
| Sequential Enc | Default Mode | 0.15 | 0.12 | 0.1 |
|  | Saliency | 0.21 | 0.17 | 0.14 |
|  | Motor | 0.17 | 0.16 | 0.11 |
|  | Visual | 0.16 | 0.13 | 0.11 |
|  | Subcortical | 0.17 | 0.15 | 0.13 |
|  | Cerebellar | 0.13 | 0.1 | 0.07 |

Our study reveals several important insights regarding the impact of architectural choices on model performance. Specifically, we observe that increasing the patch sizes within the ViT backbone leads to a modest degradation in the visual fidelity of the generated dense representations. Larger patch sizes reduce the spatial resolution of the input data, thereby limiting the model’s ability to capture fine-grained spatial details. This trade-off results in a less accurate representation of subtle, local features, which are crucial for high-quality dynamic brain mapping.

Additionally, we found that using fewer time-points in the temporal encoding significantly affects the spatiotemporal dynamics of the generated maps. When fewer time-points are included, the resulting maps exhibit more stationary behavior over time. This is because the reduced temporal granularity leads to less variation between consecutive time-points, as evidenced by a minimal mRAE between consecutive time-points, averaged across subjects. Consequently, this lack of variation results in a loss of dynamic information, diminishing the model’s ability to capture the intricate temporal fluctuations that are essential for modeling brain activity patterns over time. These findings underscore the importance of both spatial and temporal resolution in accurately capturing the dynamic behavior of brain networks and highlight the need for careful selection of patch sizes and time-points in spatiotemporal modeling tasks.

## VI. Conclusion

Our study established a new experimental paradigm for the study of brain dynamics by proposing a novel method to model both spatial and temporal dimensions from input fMRI data and generate dynamic brain maps. This approach enables us to study brain dynamics within different brain networks with a high degree of granularity. By leveraging the spatiotemporal ViT architecture and weakly supervised learning techniques, our model effectively captures the spatiotemporal variations in brain activity, even in the absence of direct ground-truth data. The model’s ability to produce smooth, temporally evolving brain maps from noisy priors offers a promising new tool for neuroimaging research, particularly in understanding complex neurological conditions such as schizophrenia. The significant differences observed between patient and control groups underscore the model’s potential for clinical applications, including differential diagnosis and personalized treatment strategies. Ultimately, this work lays the foundation for future advancements in dynamic brain mapping, providing a powerful framework for exploring brain activity in health and disease.

